# Glacier-induced upwelling shapes microbial communities in Arctic marine systems

**DOI:** 10.64898/2026.05.12.724575

**Authors:** Jenifer S. Spence, Erin M. Bertrand, Patrick L. White, Claire M. Parrott, Stephanie Waterman, David Didier, Megan E. Roberts, Andrew K. Hamilton, Maria A. Cavaco, Terry Noah, Nagissa Mahmoudi, Kurt O. Konhauser, Maya P. Bhatia

**Affiliations:** Department of Earth and Atmospheric Sciences, University of Alberta; Department. of Biology, Dalhousie University; Department of Earth, Ocean, and Atmospheric Sciences, University of British Columbia; Department of Biology, Chemistry, and Geography, Université du Québec à Rimouski; Ausuittuq Adventures, Grise Fiord, NU, Canada; Department of Earth and Planetary Sciences, McGill University

## Abstract

The Canadian Arctic Archipelago (CAA) is warming at an unprecedented rate, leading to sea ice loss and glacial retreat. Marine-terminating (tidewater) glaciers can fuel summertime marine productivity by delivering nutrient-rich deep waters via upwelling to the surface ocean. While the impact of glacier-induced upwelling has been well-studied in the context of phytoplankton and primary productivity, its effects on broader marine microbial communities remain poorly understood. We investigated how glacier-driven upwelling shapes marine microbial (bacterial and archaeal) communities across a series of sites in the CAA. At upwelling sites, the upper 50 m of the water column exhibited elevated nutrient concentrations and physical characteristics that resembled deeper waters, which were associated with differences in microbial community composition relative to non-upwelling sites. Our results indicate that upwelling influences microbial communities in surface waters in two ways. It directly introduces typically deeper-water-associated taxa into surface waters and reshapes ecological niches by enhancing nutrient supply and stimulating primary production, indirectly driving changes in microbial communities. The enrichment of *Candidatus Nitrosopumilus*, a deep water nitrifier, likely affects nitrogen cycling and raises the possibility of active nitrification in surface waters. Likewise, the increased abundance of taxa known to be associated with phytoplankton-derived organic matter in upwelling regions suggests an enhanced capacity to process organic matter generated from elevated primary productivity. Ultimately, as tidewater glaciers continue to retreat, the resulting changes in the glacially-driven upwelling regime will likely shift marine microbial communities towards assemblages adapted to less productive ecosystems, with implications for nutrient cycling in these systems.

**Importance:** Climate change has a disproportionate impact on the Arctic, with rising temperatures causing increased marine-terminating glacier retreat and changes in the marine water column structure. The consequent loss of the ability of these glaciers to upwell deep water to the surface ocean results in a reduction of nutrient delivery and mixing in these ecosystems. Previous work has highlighted the importance of marine-terminating glaciers in sustaining phytoplankton productivity during the summer season through this delivery of deep-water nutrients to the surface ocean. The impact of glacially-induced upwelling on marine bacterial and archaeal communities, however, remains underexplored. We found that in regions with glacially-driven upwelling, the surface ocean showed enrichment of phytoplankton-associated taxa and nitrifiers commonly associated with deep waters. This work underscores the role of glacially-driven upwelling in structuring both microbial communities and nutrient cycling, suggesting that glacier loss could reshape community composition and biogeochemical processes in a rapidly changing Arctic.

## Introduction

Polar environments are experiencing rapid transformations as a result of climate change (Fountain et al., 2012; Huntington et al., 2020). The Canadian Arctic Archipelago, a group of islands situated between the Arctic Ocean and Baffin Bay, is estimated to contain the largest area of land ice outside of Greenland and Antarctica (∼150,000 km^2^) (Box et al., 2018). Recent estimates indicate that glacier ice mass loss accelerated in this region in the last two decades (Gardner et al., 2011; Gardner et al., 2012; Hugonnet et al., 2021; Kochtitzky et al., 2022). Previous observations in polar regions, including the Canadian Arctic Archipelago, have demonstrated that glacial meltwater delivered into the ocean can enhance the supply of essential macronutrients (e.g., nitrogen, silicon, phosphorus) and trace metals (e.g., iron, zinc, manganese), to surface waters with important consequences for microbial food webs (Etherington et al., 2007; Bhatia et al., 2013; Hawkings et al., 2015; Juul-Pedersen et al., 2015; Hopwood et al., 2018; Kanna et al., 2018; Bhatia et al., 2021; Williams et al., 2021; White et al., 2025).

Tidewater (marine-terminating) glaciers can enrich the surface ocean with macro- and micronutrients via two mechanisms: (1) meltwater directly enriching the ocean surface (Tremblay & Gagnon, 2009; Hawkings et al., 2017; Hatton et al., 2019), and (2) subglacial discharge that drives upwelling of nutrient-rich deep water (glacially-induced upwelling) (Hawkings et al., 2016; Meire et al., 2017; Bhatia et al., 2021). The latter mechanism makes tidewater glaciers particularly important for sustaining macronutrient delivery (i.e., nitrate (NO_3_^-^), phosphate (PO_4_^4-^), silicate (SiO_4_^2-^)) into the late summer, which in turn sustains and stimulates phytoplankton growth into the autumn (Meire et al., 2017; White et al., 2025). The extent of this late-summer delivery, however, largely depends on whether the glacier terminus sits deep enough in the water column to entrain the nutrient-rich deep waters (Hopwood et al., 2018; Oliver et al., 2023). In contrast, land-terminating glaciers (glaciers whose termini end on land) deliver meltwater to the surface ocean via streams and groundwater but do not generate glacially-driven upwelling. Although meltwater can still supply macronutrients, concentrations are low relative to deep marine waters, restricting late-summer nutrient supply when considered alone (Hopwood et al., 2018; Hopwood et al., 2020). Consequently, marine regions influenced primarily by land-terminating glaciers tend to be less productive than those influenced by marine-terminating glaciers (Meire et al., 2017).

While glacier-driven upwelling has been shown to shape eukaryotic phytoplankton communities and primary production (Halbach et al., 2019; Meire et al., 2023; Roberts et al., 2024; White et al., 2025), its effects on the bacterial and archaeal communities remain comparatively poorly constrained. Nutrient delivery via glacially-driven upwelling can increase the production and export of phytoplankton-derived organic matter that supports and structures heterotrophic microbial communities. For instance, dominant Arctic phytoplankton taxa, including bloom-forming diatoms, *Micromonas* spp., and *Phaeocystis* (Monier et al., 2015; Joli et al., 2018; Kalenitchenko et al., 2019), release lysates and metabolic by-products which fuel heterotrophic and mixotrophic microbial activity (Schoemann et al., 2005; Buchan et al., 2014; Han et al., 2024). Dominant Arctic marine hetero- and mixotrophic microorganisms, including *Polaribacter* (Flavobacteriaceae) and *Sulfitobacter* (Rhodobacteriaceae), have been previously linked to phytoplankton blooms due to their ability to degrade high molecular weight organic compounds (Malmstrom et al., 2007; Ghiglione et al., 2012; Moran et al., 2012; Teeling et al., 2012; Pedrós-Alió et al., 2015). In addition to organic matter supply, glacially-driven upwelling alters the availability of nitrogen, particularly by increasing NO_3_^-^ inputs from deeper waters (Meire et al., 2017). As a result, the microbial community may shift toward taxa that favour “fresh” nitrogen (e.g., NO_3_^-^ delivered from deeper waters,) versus those adapted to “regenerated” nitrogen (e.g., nitrogen species cycled within the euphotic zone, such as amino acids and urea) (Baer et al., 2017). Together, these observations suggest that glacier-induced upwelling can structure the microbial community through both bottom-up stimulation of organic carbon supply and shifts in nutrient regime.

In addition to changes in carbon and nutrient availability, tidewater glaciers can also influence microbial communities via physical processes. One such mechanism is through the delivery of glacially-derived taxa to the surface ocean via meltwater fluxes. Indeed, studies examining bacterial distributions along the glacier ice-to-ocean continuum have documented shared taxa across supraglacial, subglacial, proglacial river/stream, and marine environments (Comte et al., 2018; Thomas et al., 2020). A second physical influence may be through the delivery of deep-water taxa to the surface ocean via glacially-induced upwelling of deep seawater to the surface ocean. The presence of deep water taxa in the surface ocean has been previously observed across the global ocean and has been attributed to vertical mixing (Alves Junior et al., 2015; Wenley et al., 2021). The enhanced vertical mixing driven by glacially-induced upwelling therefore represents a plausible mechanism by which tidewater glaciers can restructure surface microbial communities, although this process has not yet been explicitly investigated in tidewater glacier influenced systems.

In this study, we investigated the influence of tidewater glacier-induced upwelling on marine microbial communities in the CAA. Specifically, we: (1) assessed whether glacially-induced upwelling influences the microbial community composition; and (2) determined which ecological, environmental, and physical factors may drive these patterns. To do so, we first identified the presence of active tidewater glacier-driven upwelling across Jones Sound and Nares Strait in Inuit Nunangat (Nunavut, Canada) via examination of physical and geochemical water properties. We then applied 16S rRNA gene amplicon sequencing to investigate bacteria and archaeal (here-on referred to as microbial) communities at sites with and without active glacially-induced upwelling. Overall, this comparison identified consistent trends between tidewater glacier-driven upwelling and increased relative abundances of putative phytoplankton-associated taxa and deep-dwelling nitrifiers in the surface ocean. Our results suggest that with changes in the glacially-driven upwelling regime, there may be a shift in the microbial community towards taxa better adapted for more oligotrophic conditions, which could lead to a restructuring of surface ocean biogeochemical cycling.

## Methods

### Glacier Inputs and Oceanographic Setting

Water column sampling and in situ measurements were conducted during August 2021 and August and September 2022 in Jones Sound and Talbot Inlet (Nunavut, Canada) (Figure 1). Jones Sound is a waterway situated between Ellesmere and Devon Islands. Each summer, Jones Sound receives glacial meltwater draining the Sydkap and Manson Ice Fields on Ellesmere Island to the north, and Devon Ice Cap on Devon Island to the south. Talbot Inlet is situated on the southeastern side of Ellesmere Island, near the southern reaches of Nares Strait, and contains several tidewater and land-terminating glaciers draining the Prince of Wales (POW) Icefield. The two largest outlet glaciers, Trinity and Wykeham glaciers, are located at the head of Talbot Inlet and have undergone a significant increase in calving, velocity, and iceberg discharge over the last 15 years (Dalton et al., 2019; Dalton et al., 2025). See Bhatia et al. (2021), Roberts et al. (2024), and White et al. (2025) for further oceanographic context for this region.

**Figure 1.**
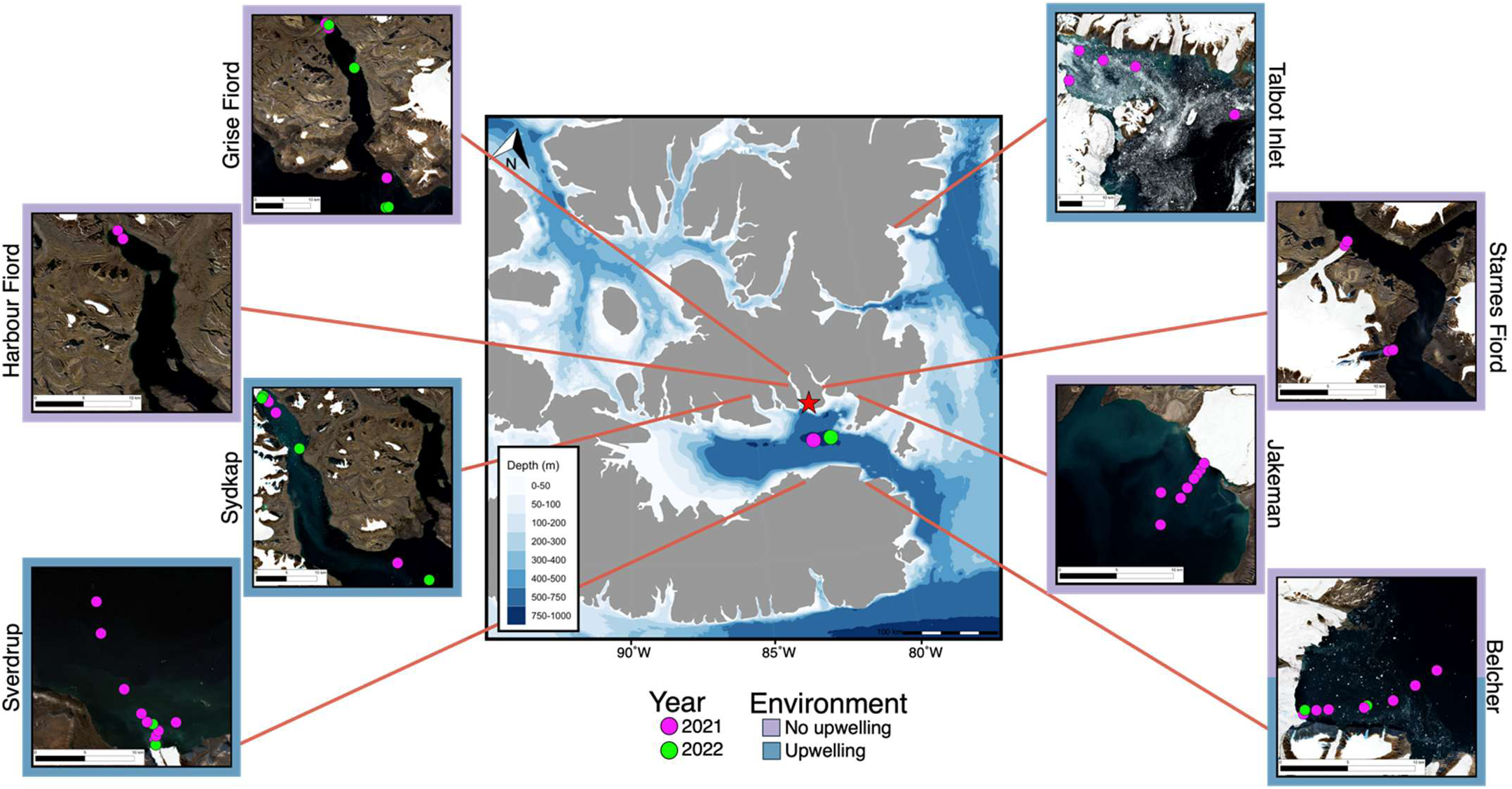
Map of stations across Jones Sound and Nares Strait in Inuit Nunangat (Nunavut, Canada), with the community of Ausuittuq denoted by the red star. Insets show locations of bottle stations at 10 sites spanning 2021 and 2022 (coloured circles). Six sites had active glacier-induced upwelling (Belcher (2022), Sydkap (2021, 2022), Sverdrup (2021, 2022), Talbot Inlet (2021)). Seven sites had no glacier-induced upwelling (Belcher (2021), Jakeman (2021), Starnes.1 (2021), Starnes.2 (2021), Harbour Fiord (2021), Grise Fiord (2021, 2022). Samples collected from Jones Sound (2021, 2022) and Nares Strait (2021) were classified as open water. Inset maps were generated in QGIS (v.3.44.6) using Copernicus sentinel data from august 2019 processed by Sentinel Hub. Centre map was generated using the R package ggOceanMaps where land polygons are made with Natural Earth and bathymetry is created with IBCAO data (v. 4.0) (Jakobsson et al., 2020; Vihtakari, 2022). See Figure S1 for a detailed station map.

In total, six sites with tidewater glaciers and three sites with no tidewater glaciers were sampled. Sites with tidewater glaciers were Belcher, Sverdrup and Sydkap (sampled in 2021 and 2022), and Jakeman, Starnes.1, and Talbot Inlet (sampled in 2021). The sites with no tidewater glaciers present were Grise Fiord (sampled in 2021 and 2022), and Harbour Fiord and Starnes.2 (sampled in 2021). Additionally, open water samples were collected in Nares Strait (2021) and Jones Sound (2021, 2022). Sampling was performed aboard the sailing yacht (*S/Y) Vagabond* (2021) and community owned/operated vessels (*Ausuittuq Adventures*) from Grise Fiord (Ausuittuq) (2021, 2022). The *S/Y Vagabond* was used to sample tidewater glacierized sites in Talbot Inlet and on the southern coast of Jones Sound, and the community vessels were used to sample the northern coast of Jones Sound. Additionally, the *S/Y Vagabond* was used to sample the open water stations in Nares Strait in 2021 (i.e., Talbot_Inlet_1), and Jones Sound in both 2021 and 2022.

### Site Classifications

Not every site with a tidewater glacier present may not be actively upwelling nutrient-rich deepwater at the time of sampling. Thus, it is important to make the distinction between the presence of tidewater glaciers and the presence of active tidewater-glacier induced upwelling. For this reason, an upwelling indicator was designed to determine whether active glacier-induced upwelling was occurring at the time of sampling at Belcher, Sverdrup, Sydkap, and Talbot Inlet (Parrott et al., Submitted Oct. 2025). These sites were selected for their capacity for glacier-induced upwelling, as their grounding lines sit below the nutricline (Bhatia et al., 2021; Roberts et al., 2024; White et al., 2025). The upwelling indicator was determined based on in situ measurements of turbidity, dissolved oxygen (O_2_), and temperature anomalies, as well as a measure of isopycnal uplift. To calculate these anomalies, the vertical profile of each variable at each station, within 10 m above and below the 1026 kg m^-3^ isopycnal, was subtracted from the corresponding mean profile at open water reference stations in Jones Sound or Nares Strait for the relevant sampling year. The 1026 kg m^-3^ isopycnal was used as it has been shown to be an indicator of upwelling (Bhatia et al., 2021). The isopycnal uplift was computed by taking the change in depth of the 1026 kg m^-3^ isopycnal between the station in question and the outermost station of the transect, except at Sverdrup Glacier where the inner stations are compared to the outermost station inside the basin sill (Figures S2-S6). Negative anomalies of temperature and dissolved O_2_ were considered as an indication of active upwelling. Positive values of isopycnals uplift and positive anomalies of turbidity were considered to be an indication of active glacier-induced upwelling. Variables were median-centered and scaled by their interquartile range (IQR = 1). The upwelling indicator for each station was calculated as the sum of the scaled variables. Typically, the impacts of glacially-induced upwelling are observed within 4 to 5 km of a glacier’s terminus in the Canadian Arctic Archipelago (Bhatia et al., 2021; Williams et al., 2021), therefore this distance was used as an outer boundary in computing the 1026 kg m^-3^ isopycnal uplift to better constrain upwelling as glacier-induced.

Based on the upwelling indicator, samples collected from the upper 50 m from Belcher Glacier in 2022, Sverdrup Glacier in 2021 and 2022, Sydkap Glacier in 2021 and 2022, and Talbot Inlet in 2021 were characterized as having active glacier-induced upwelling at the time of sampling, referred herein as ‘Upwelling’ (Figure 1). Samples characterized as having no active glacier-induced upwelling at the time of sampling (‘No Upwelling’) in the upper 50 m of the water column were Belcher Glacier in 2021, Grise Fiord in 2021 and 2022, Harbour Fiord in 2021, Jakeman Glacier in 2021 2021, as well as Starnes.1 and Starnes.2 in 2021. Samples collected from the upper 50 m of open water sites were classified as ‘Open Water’. Due to similar physical and geochemical signatures, all water collected below 50 m, regardless of site, were classified as ‘Below 50 m’.

### Water Column Sampling

Water samples were collected for analysis to determine microbial community composition and geochemical characteristics. A total of 51 Upwelling samples, 44 No Upwelling samples, 13 Open Water samples, and 37 samples from below 50 m depth. Sampling was conducted at each station and depth as described in White et al. (2025), with CTD-casts conducted wherever water samples were collected. Summaries of sample metadata and section plots of physical variables are in supplementary materials (Suppl. Table 1, Suppl. Figures 1-5).

In situ measurements of physical and geochemical water column properties were collected using a RBR maestro^3^ multichannel logger (RBR Ltd.) equipped with sensors for conductivity, temperature, pressure, dissolved O_2_, chlorophyll *a* fluorescence (Chl*a*), and turbidity. Data collection proceeded as described by Bhatia et al. (2021). Sensor measurements were processed using Ruskin’s MATLAB toolbox, RSKtools (https://rbr-global.com/products/software) and MATLAB (v.2022b; Mathworks, Natick, MA, USA), and data was averaged in 0.5 m depth bins. Buoyancy (N2; herein referred to as stratification) was calculated with the GSW toolbox using a 10-point (5 m) running average (McDougall & Barker, 2011).

Water column samples were collected from up to 3 different depths in the water column. Specifically, samples were collected from the surface environment (5-8 m depth), the subsurface chlorophyll maximum (SCM) (18-50 m depth), and deep in the water column (> 50 m depth). If there was no observable peak of Chl*a* in the SCM, water was collected at 20 m depth. Bottle samples were collected using a 10 L (Model 1080, non-metallic) GO-FLO on the *S/Y Vagabond*. On the community vessels, either a 10 L (Model 1080, non-metallic) GO-FLO, 1.7 L Niskin Model 1010, or 2.5 L GO-FLO Model 1080 (non-metallic) was used to collect bottle samples. All GO-FLO and Niskin were cleaned according to trace-metal clean procedures in the laboratory prior to the field (Cutter & Bruland, 2012; Bhatia et al., 2021).

Water samples for macronutrient analysis (i.e., concentrations of nitrate [NO_3_^-^]; ammonia [NH_3_]; phosphate [PO_4_^3-^]; silicate [SiO_4_^4-^]) were collected by filling a 60 mL rubber syringe directly from the nozzle of the GO-FLO or Niskin bottle and filtering through a 0.22 µM polyethersulfone (PES) syringe filter into a clean 15 mL centrifuge tube (rinsed 3x with filtered sample water). Macronutrient samples were frozen upright immediately following collection aboard the *S/Y Vagabond* or kept in a dark cooler on the community boat until return to Grise Fiord where samples were subsequently frozen at -20 °C within 10 hours of sampling. Macronutrient concentrations were analyzed on a Skalar SAN++ Continuous Flow Nutrient Analyzer at the CERC OCEAN Laboratory (Dalhousie University). The limits of detection for NO_3_^-^, NH_3_, PO_4_^3-^, and SiO_4_^4-^ concentrations were 0.01, 0.05, 0.01, and 0.05 µM, respectively. The limits of quantification for NO_3_^-^, NH_3_, PO_4_^3-^, and SiO_4_^4-^ concentrations were 0.16, 0.19, 0.04, and 0.24 µM, respectively. For any values that fell below the limits of detection, the raw instrument values were used instead (Antweiler, 2015).

We filtered 1-2 L of water for 16S rRNA gene amplicon analysis through 0.22 µM Sterivex™ filters using a peristaltic pump, at a rate of 50-60 mL/min). On the *S/Y Vagabond*, water samples for DNA were filtered immediately after collection and then flash frozen at -80 °C. DNA samples collected aboard community vessels were stored in coolers in the dark until return to Grise Fiord, where they were filtered and frozen at -20 °C within 10 hours of water collection.

### Nucleic Acid Extraction and Amplicon Sequencing

Total DNA was extracted from the Sterivex™ filters using the DNeasy PowerWater Sterivex Kit (Qiagen). All steps in the extraction kit protocol were followed, apart from the heated incubation step, which was extended to a 1-hour incubation at 70 °C to maximize DNA yield recovered. Following extraction, DNA concentrations were determined using the QuBit dsDNA HS Assay Kit™ on a Qubit 3.0. Extracted DNA was used to characterize the microbial community composition using 16S rRNA gene sequencing. The V4V5 hypervariable region (515F/926R) of the 16S rRNA gene was targeted, with primers containing a six-base index sequence for sample multiplexing (Bartram et al., 2011; Quince et al., 2011; Parada et al., 2016). The PCR mix and amplicon sequencing were prepared and conducted as described in Roberts *et al*. (2024).

### Sequencing and Statistical Analyses

Amplicons were analyzed using QIIME2 v.2021.4, as described by Roberts et al. (2024). Briefly, amplicon sequence variants (ASVs), were classified using the SILVA reference database (v.138, 2019), referred herein as the microbial community (bacteria, archaea) (Klindworth et al., 2013; Bolyen et al., 2019). ASVs assigned to chloroplasts and mitochondria were filtered from the ASV dataset. The resulting curated ASV table was used for downstream statistical analysis and visualization in R (v.4.1.2). Samples with less than 2000 reads were dropped from the microbial community analysis. Relative abundance for community composition analysis was calculated from raw reads for taxonomic bar plots. Sequence data was Hellinger transformed prior to ordination and statistical testing by non-metric multidimensional scaling (NMDS). A pairwise permutational multivariate analysis of variance (PERMANOVA) was calculated to test for differences among site classifications for both the physical (turbidity, temperature, salinity, Chl*a*, stratification) and geochemical (dissolved O_2_ and macronutrient concentration) variables as well as the microbial community, with the p-value corrected for multiple comparisons using the Holm adjustment method (Hervé & Hervé, 2020). A principal component analysis (PCA) was used to assess spatial variability in the physical and geochemical properties of the water column, with a focus on the presence or absence of glacially-driven upwelling. Data was scaled around the mean prior to calculating the Euclidean distances. For both PCA and NMDS visualizations, only ellipses indicating significant differences between the depth ranges (above 50 m depth versus below 50 m depth) and Upwelling versus No Upwelling groupings are shown for ease of interpretation. All adjusted p-values from PERMANOVA analysis are summarized in Suppl. Table 2.

A distance-based redundancy analysis (db-RDA) was used to quantify the variance in the microbial community composition in a Bray-Curtis matrix explained by normalized physical and geochemical characteristics in the upper 50 m of the water column. Physical and geochemical variables were selected as significant using the forwards-backwards stepwise db-RDA and checked for multicollinearity (>10) using the Vegan package for R (Oksanen et al., 2013). Of note, the [NO_2_^-^] and stratification variables were not significant and therefore excluded, while [SiO_4_^4-^] was removed due to collinearity with [NO_3_^-^]. Samples that lacked one or multiple water physical or geochemical parameters were excluded from analysis for the db-RDA (n = 2; Supp. Table 1).

To identify taxa significantly associated with active glacier-induced upwelling, a Multinomial Species Classification Method (CLAM) was used (Chazdon et al., 2011). To better capture the impacts of glacier-induced upwelling on microbial community composition, only water collected within 5 km of the glacier terminus or shoreline was considered for this analysis. Taxa associated with environments with active upwelling (n=35) and no active upwelling (n=28) were determined. Each ASV was classified into a category: taxa associated with upwelling habitats, taxa associated with habitats lacking upwelling, generalist (i.e., no statistically significant difference in read counts between habitats), or too rare. A threshold was established to discern between ASVs that were present at sites with upwelling due to the direct delivery from deeper waters to the surface, or the generation of new ecological niches. Taxa associated with upwelling environments that exhibited an average relative abundance of ≥1.5% below 50 m depth were interpreted as being present in the upper 50 m as a result of being transported with upwelled deep water. All other upwelling-associated taxa were considered to be present due to the indirect impacts of upwelling through the generation of new niches, such as an increase in organic matter production due to enhanced primary productivity or nutrient availabilities in the upwelling environments. In visualizations, only the ASVs that had an average relative abundance >0.5% within each habitat were displayed.

## Results

### Influence of upwelling on water column physical and chemical variability

There was a clear separation in physicochemical properties associated with depth (Figure 2a; PERMANOVA p-value < 0.05). In general, waters below 50 m depth were characterized by elevated salinity, [PO_4_^3-^], [SiO_4_], and [NO_3_^-^], while samples from the upper 50 m of the water column were characterized by elevated dissolved O_2_, stratification, temperature, and Chl*a*. Within the upper 50 m, water column properties varied according to the influence of upwelling (Figure 2b). Stations from sites with active upwelling were significantly distinct in their geochemical and physical variables from those without upwelling influence (PERMANOVA p-value < 0.05). This finding is not surprising given the upwelling indicator was designed to classify sites based on upwelling signatures from some of these same physiochemical properties. Waters collected from upwelling sites were generally characterized by elevated macronutrient concentrations, salinity, and Chl*a* fluorescence. An exception to this were some samples from Sydkap, which were characterized by elevated NH_3_ and turbidity (Figure S6). These signatures likely reflect riverine input rather than upwelling, as supported by satellite imagery (Figure S7). In contrast, samples classified as No Upwelling and Open Water were typically characterized by increased temperature and stronger stratification.

**Figure 2.**
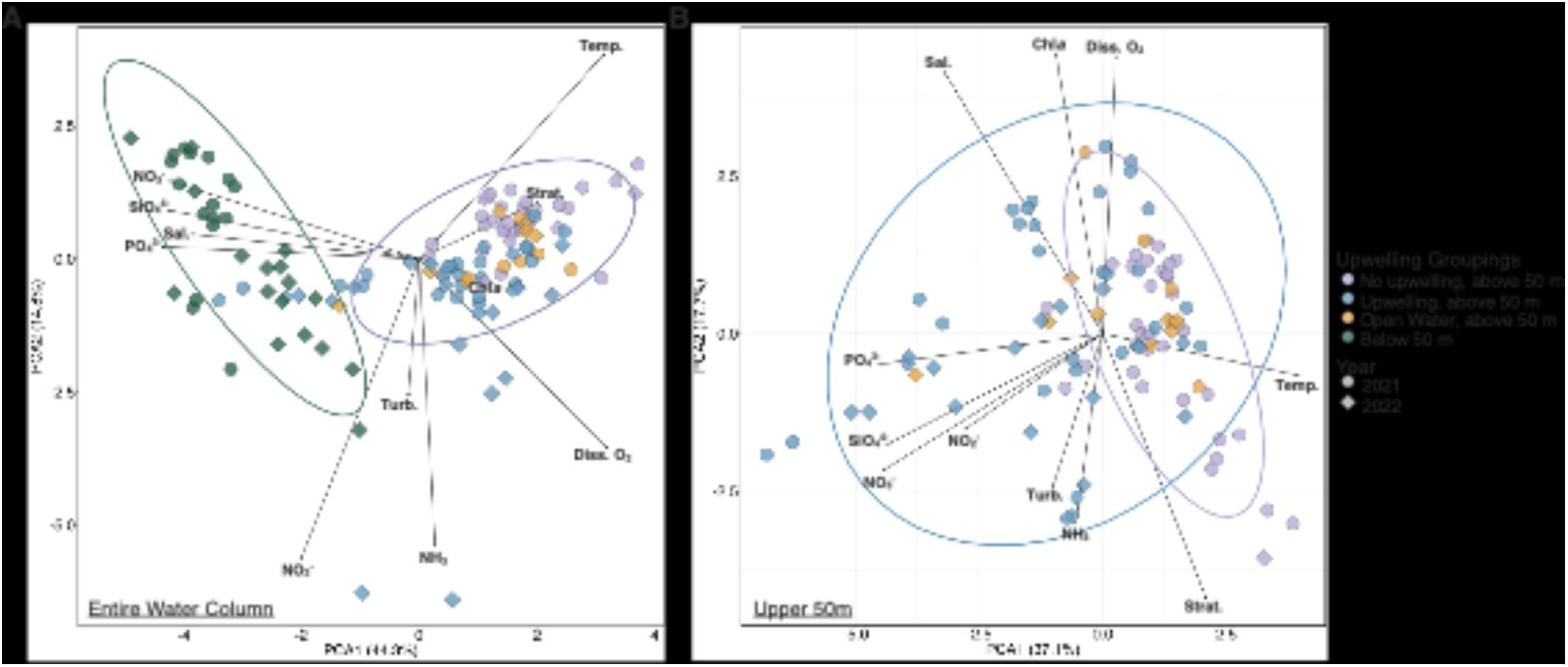
Principal component analysis (PCA) using a Euclidean distance matrix of physical (strat., turb., temp., diss. O_2_, Chl*a*) and chemical (NO_3_^-^, NO_2_^-^, NH_3_, PO_4_^3-^, SiO_4_^4-^) variables of the (A) entire water column (green ellipse = below 50 m, grey ellipse = 50 m and above) and (B) upper 50 m of the water column (blue ellipse = upwelling, upper 50 m within 5 km, purple ellipse = no upwelling, upper 50 m within 5 km). Ellipses represent 95% confidence intervals and are significantly different from one another (PERMANOVA p-value < 0.05). Shapes represent the years sampled, and colours indicate the sample classification (see methods). Strat. = stratification; turb. = turbidity; temp. = temperature; diss. O_2_ = dissolved oxygen; Chl*a* = chlorophyll *a*. Percentage values on axis labels represent the total variance in the dataset explained by that principal component.

### Microbial community composition

Of the 6,727 ASVs identified in the entire water column, 4,419 of these microbial ASVs were observed in the upper 50 m of the water column. The overall community composition remained broadly consistent across sites and depths, some taxa showed marked shifts in relative abundance between site classifications (Figure 3). *Polaribacter* dominated the community, with a mean relative abundance of 29%, with values ranging from 12% in waters below 50 m depth to 39% in upwelling-induced surface waters. Taxa with an average relative abundance of <1% collectively constituted a substantial fraction of the community, ranging from a minimum of 15% in open surface waters to a maximum of 37% in water below 50 m depth (mean: 22%). This distribution pattern is reflected in community diversity, with the highest Shannon Index (SI) observed in waters below 50 m depth (Figure S8).

**Figure 3.**
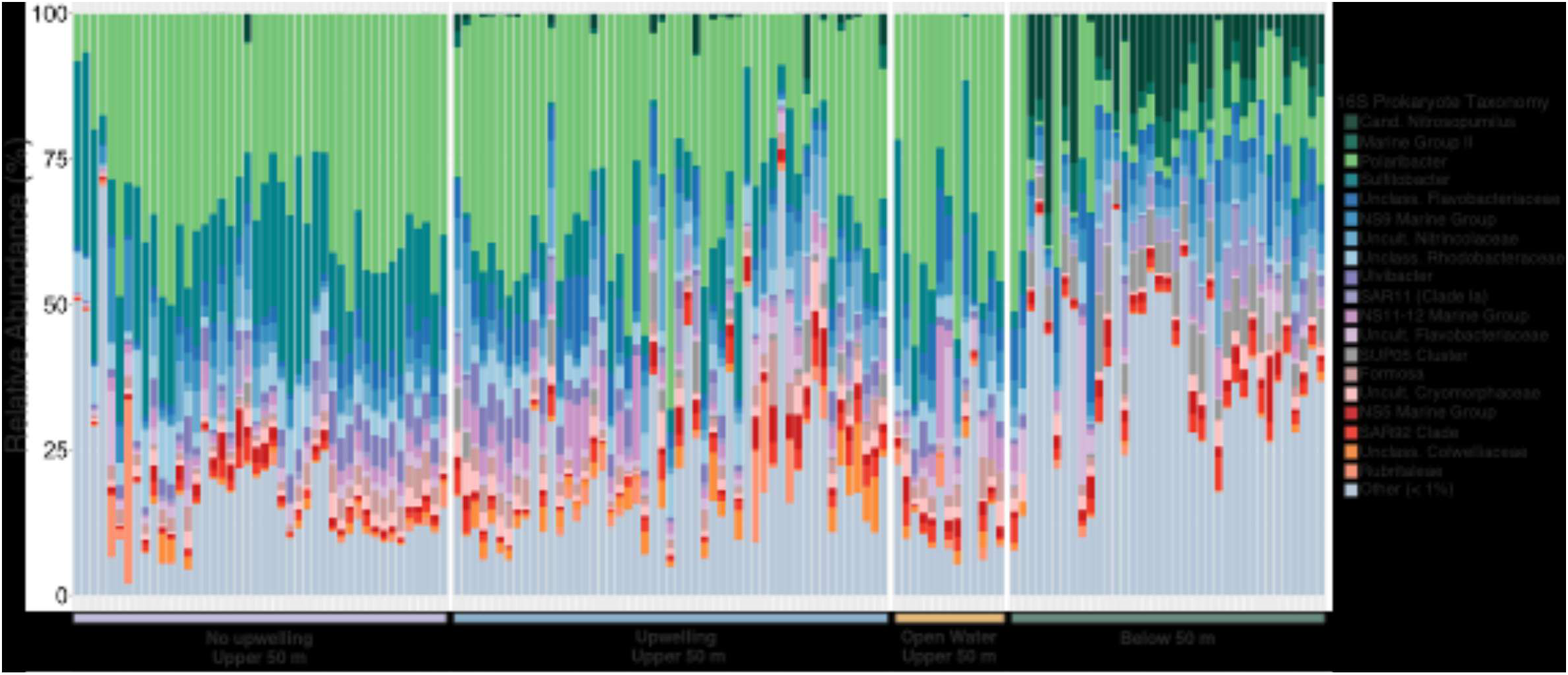
Average relative abundance of microbial taxonomic groups determined by 16S rRNA amplicon sequencing in the entire sampled water column (depth range: 5-500 m). Genera found at less than 1% average relative abundance were categorized as “Other” (<1%). Samples are grouped according to their upwelling classification, indicated by the coloured bar.

Water below 50 m was enriched in archaeal taxa across all sites, most notably *Candidatus Nitrosopumilus* (10%) and *Marine Group II* (3.6%) (Figure 3). In contrast, dominant surface water bacteria declined with depth: both *Polaribacter* and *Sulfitobacter* decreased to 12% and 1.2%, respectively, with *Sulfitobacter* being nearly undetectable in over half of the deep-water samples. Other deep-water taxa included *SUP05* (5.9%) and *SAR11* (4.3%). Conversely, the upper 50 m of the water column was dominated by *Polaribacter* (35%) and *Sulfitobacter* (12%), with the latter exhibiting relatively higher relative abundances at sites lacking upwelling (19%) compared to upwelling sites (7.4%). Both *RS62 Marine Group* and *Candidatus Aquiluna* had increased relative abundance in the upper 50 m relative to deep water. However, their presence appeared to be more site specific: *RS62* was largely confined to stations at the Sydkap and Jakeman sites, whereas *Cand*. *Aquiluna* showed variable relative abundance across the No Upwelling (1.7%), Upwelling (0.8%), and Open Water (1.8%) surface sites (Figure S9).

In the upper 50 m, sites influenced by glacially-induced upwelling had an overall increased relative abundance of unclassified *Colwelliaceae* (2.3%) compared to sites without upwelling (0.7%). Additionally, stations in upwelling regions showed an enrichment in taxa such as *Cand. Nitrosopumilus*, *SUP05*, and *SAR11*, with maximum relative abundances of 11%, 8.9%, and 10%, respectively (Figure 3). Notably, three stations at sites with no active glacially-induced upwelling also showed enrichment in some, or all, of these taxa. A station along the Jakeman site (Jakeman_44b; Figure S1) exhibited an enrichment in *Cand. Nitrosopumilus* (up to 4.7%)*, SAR11* (up to 6.7%), and *SUP05 Cluster* (up to 3.3%) at either or both sampled depths (5 m and 20 m). All stations at both Starnes sites were enriched in *SAR11* (up to 8.3%).

### Influence of tidewater glaciers on microbial community composition

A NMDS analysis was used to evaluate dissimilarities in microbial community composition throughout the water column and between sites with and without glacier-induced upwelling. A significant difference was observed between communities above and below 50 m depth (PERMANOVA p-value < 0.05; Figure 4a, Table S2). Similarly, in the upper 50 m of the water column there was a clear separation between sites based on the presence or absence of upwelling, as well as between the No Upwelling and Open Water (above 50 m) groups (PERMANOVA p-value < 0.05; Figure 4b, Table S2).

**Figure 4.**
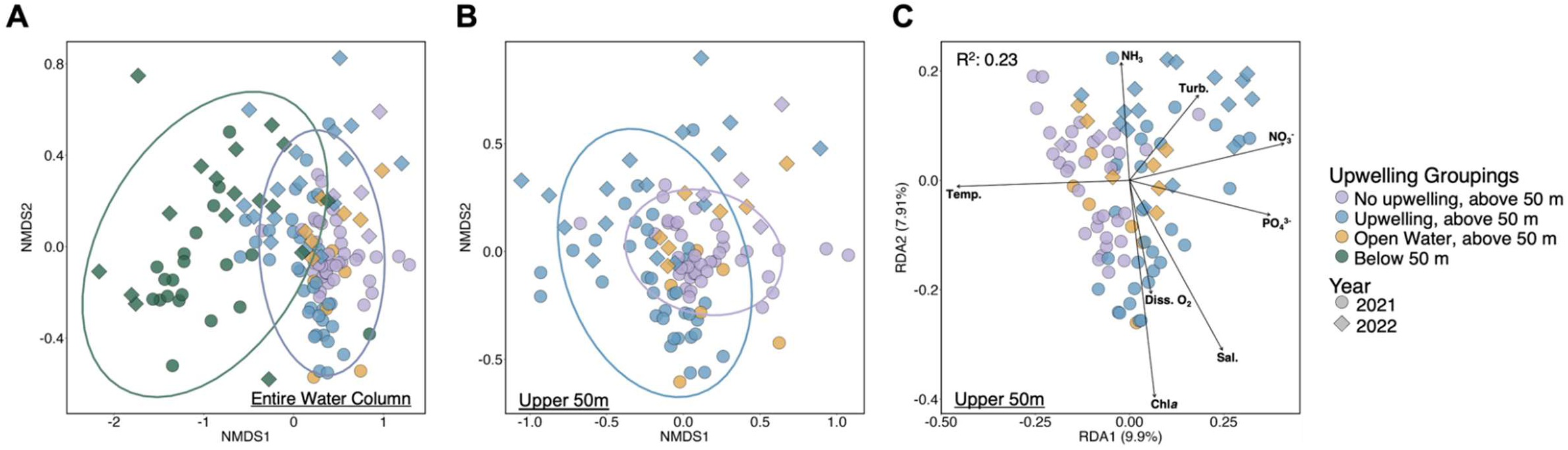
Microbial community composition utilizing a non-metric multidimensional scaling plot (NMDS) ordination of the (A) entire water column (stress = 0.11; green ellipse = below 50 m, grey ellipse = 50 m and above) and (B) upper 50 m of the water column (stress = 0.18; blue ellipse = upwelling, upper 50 m within 5 km, purple ellipse = no upwelling, upper 50 m within 5 km). (C) Distance-based redundancy analysis (dbRDA) of the microbial community composition in the upper 50 m of the water column with phosphate (PO_4_^3-^), nitrate (NO_3_^-^), ammonia (NH_3_), salinity (Sal.), turbidity (Turb.), temperature (Temp.), and dissolved oxygen (Diss. O_2_ (μM)). Adjusted R^2^ represents the proportion of variation that is explained by the significant environmental predictors (adj. R^2^ = 23%). Percentage value on axis labels represent the proportion of model-explained variance. Vectors indicate significant physical and chemical predictors (P < 0.05) as selected by forwards-backwards selection. Shapes represent the years sampled, and colour indicates the sample classification (see methods). Ellipses represent 95% confidence intervals and are significantly different from one another (PERMANOVA p-value < 0.05).

A dbRDA was used to assess the relationship between environmental parameters and community variation. Physical and geochemical variables together explained 23.1% of the variance in the microbial community composition (Figure 4c). Significant predictors (p-value < 0.05) included [NO_3_^-^], [NH_3_], [PO_4_^3-^], [O_2_], salinity, Chl*a*, temperature, and turbidity. In particular, microbial communities at sites with upwelling were significantly associated with increased turbidity, [NO_3_^-^], [PO_4_^3-^], salinity, and [O_2_] (Figure 4c), whereas No Upwelling communities were influenced by increased temperature. A notable exception was observed at the No Upwelling site, Jakeman Glacier, where both depths from the same station (5 and 20 m) clustered with the Upwelling grouping and were associated with increased turbidity (Jakeman_44b; Table S1, Figure S11c). In contrast, all stations at the Starnes.1 and Starnes.2 sites exhibited an increased relative abundance of *SAR11* relative to other sites without active glacially-induced upwelling, but did not cluster closely with the Upwelling groupings. Instead, these stations were associated with elevated NH_3_ concentrations (Figure S11c).

A pairwise CLAM test was used to identify surface ocean ASVs associated with active glacier-induced upwelling versus non-glacier-induced upwelling habitats. The analysis was restricted to samples from sites classified as either Upwelling or No Upwelling, collected from the upper 50 m of the water column and located within 5 km of the glacier terminus or shoreline. Out of the resulting 3,031 ASVs present across this sample subset, 711 ASVs and 244 ASVs were associated with Upwelling and No Upwelling environments, respectively. Among these, 32 Upwelling- and 10 No Upwelling-associated ASVs had mean relative abundances >0.5% within their respective habitats. Taxa associated with Upwelling environments included ASVs affiliated with unclassified *Colwelliaceae, Colwellia, Polaribacter*, and members of the Nitrincolaceae family. Apart from the *Tychonema* ASV and a single *NS3a Marine Group* ASV, the remainder of the Upwelling-associated ASVs were present below 50 m depth, ranging from 0.01-28% average relative abundance in these deeper waters. Interestingly, the cyanobacteria *Tychonema* was observed exclusively in the water column at a single Sydkap (2022) station, where it reached a relative abundance of 39%. In contrast, taxa associated with environments that had no active glacially-induced upwelling were mostly absent below 50 m depth (Figure 5b). Exceptions included ASVs belonging to the *Sulfitobacter*, *Fluviicola*, *Burkholderia-Caballeronia-Parabvurkholderia* genera, which ranged from 0.04 to 2.8% in the deep waters. Some ASVs were represented by multiple ASVs with differing habitat affinities; for example, separate ASVs from the *NS3a Marine Group* genus were identified as being associated with both Upwelling and No Upwelling environments.

**Figure 5.**
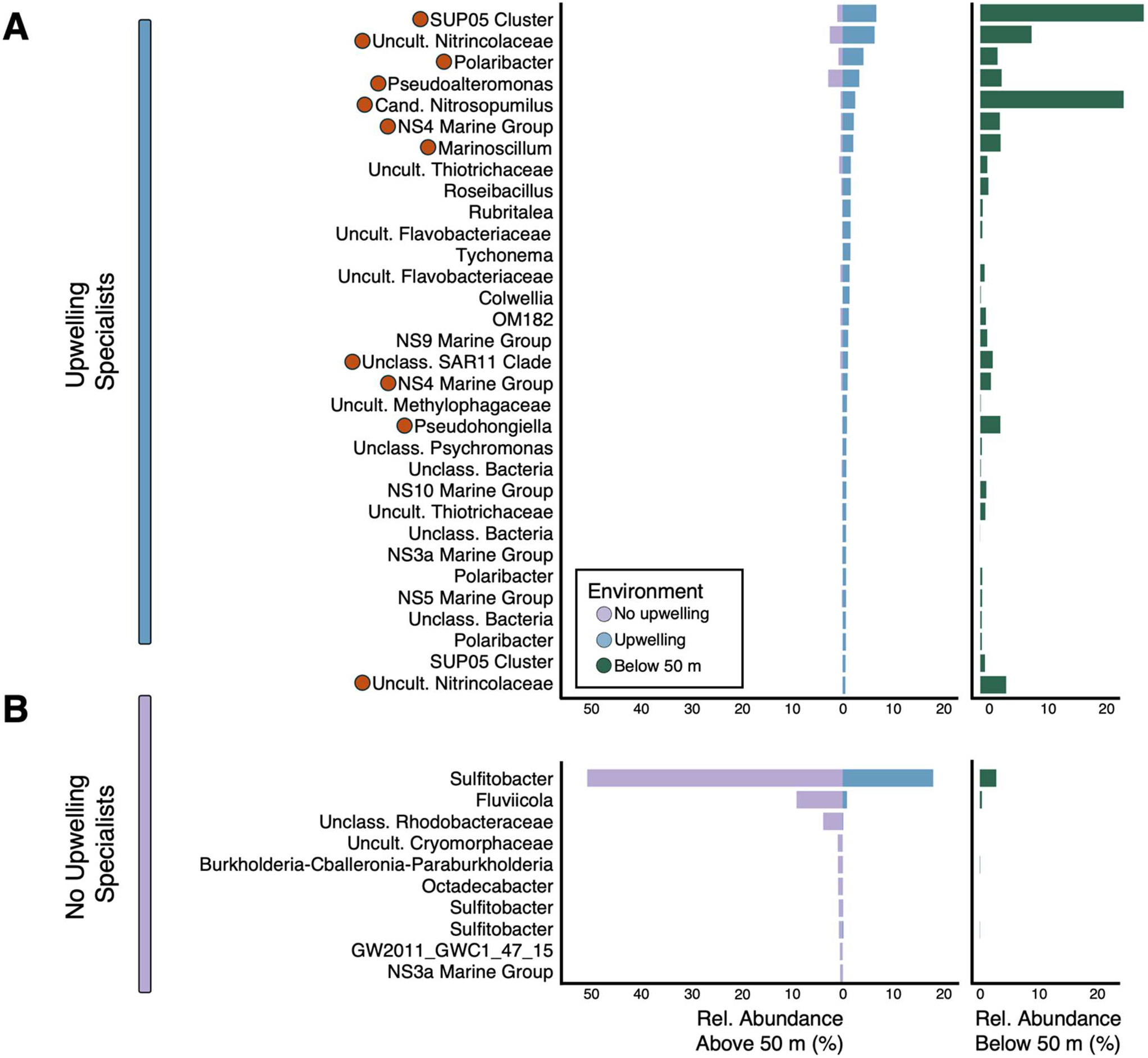
Taxa in the upper 50 m of sites associated with (A) upwelling, and (B) no upwelling, identified through the multinomial species classification method (CLAM). Purple bars represent the average relative abundance for ASVs associated with no upwelling environments in the upper 50 m, and blue bars represent the average relative abundance of ASVs associated with upwelling environments in the upper 50 m. Green bars represent the average relative abundance of taxa below 50 m. Orange circles indicate ASVs that were predicted to be present in the upper 50 m as a result of being upwelled along with deep waters based on them exhibiting an average relative abundance of ≥1.5% below 50 m depth. Cutoff threshold for visualization of CLAM-identified environment-associated ASVs are those with an average relative abundance greater than 0.5%.

## Discussion

### Influence of glacier-driven upwelling on the microbial community through enhanced productivity

There has been a recent focus on characterizing the marine microbial community in the Arctic, resulting in a growing list of commonly observed taxa (Alonso-Sáez et al., 2012; Ghiglione et al., 2012; Wilson et al., 2017; Thiele et al., 2022; Thiele et al., 2023). The dominant genera observed in this study, *Polaribacter* and *Sulfitobacter*, have been previously reported in the Arctic during summer months, consistent with the concept of a seasonal, pan-Arctic circumpolar biome (Kalenitchenko et al., 2019). While the relative contributions of *Polaribacter* and *Sulfitobacter* to community composition were variable across the sampled sites, these consistently remained the two most abundant genera in the upper 50 m of the water column. This persistence may be due to their capacity to use phytoplankton-derived macromolecules (Teeling et al., 2012; Buchan et al., 2014; Francis et al., 2021; Buchan et al., 2025), which are enriched in surface waters. In particular, *Polaribacter* spp. are a well-studied polar taxon that have been linked to increased inputs of phytoplankton-derived organic matter and often used as an indicator of such conditions (Underwood et al., 2019). The predominance of the *Polaribacter* and *Sulfitobacter* genera in the upper 50 m of the water column suggests that their distributions are not strongly impacted by glacier-driven upwelling. Instead, their prevalence likely reflects the parallel ubiquity of the diatom *Chaetoceros* in the study area (White et al., 2025), whose organic matter both genera have been shown to degrade and utilize (Bennke et al., 2013; Crenn et al., 2018; Liu et al., 2020; Tisserand et al., 2020). Although *Polaribacter* and *Sulfitobacter* were widespread across all sites, the presence of different ASVs associated with both upwelling and non-upwelling conditions suggests fine-scale niche differentiation at the species level. This suggests that these species are able to take advantage of varying ecophysiological niches, indicating an indirect impact of upwelling at the species level.

We found that glacier-induced upwelling increased the concentration of nutrients in the surface waters, such as NO_3_^-^. Increased [NO_3_^-^] in the surface ocean at regions with tidewater glaciers has been previously linked with stimulating late summer primary productivity. Further, dissolved O_2_ and Chl*a* were found to be significant predictors of microbial community structure, and were elevated in upwelling regions, suggesting a link between phytoplankton productivity and microbial community composition. Taxa observed in environments with active glacially-induced upwelling in this study include *Colwellia*, uncultured *Nitrincolaceae*, and NS (North Sea) lineages (*NS3a*, *NS4*, *NS5*, *NS9*, *NS10*). These taxa have all been previously reported in association with early-stage phytoplankton blooms in the Arctic, and are thought to be well adapted to exploit organic matter-rich conditions due to their capacity to degrade complex carbohydrates derived from phytoplankton biomass (Luria et al., 2017; Wietz et al., 2021; Thiele et al., 2022; Thiele et al., 2023). Given that sampling occurred late in the melt season, when waters would normally be nutrient-limited, the persistence of these bloom-associated microbial taxa at sites with active upwelling likely reflects a continued supply of phytoplankton-derived organic matter sustained from glacier-driven nutrient delivery.

Regions lacking upwelling were instead characterized by reduced macronutrient concentrations, increased stratification, and increased temperature. Previous work has linked these marine conditions to reduced primary productivity (Roberts et al., 2024; White et al., 2025). The identification of ASVs belonging to the *Sulfitobacter* and *Octadecabacter* genera, both considered to be copiotrophs, at sites with no active glacially-induced upwelling was thus unexpected (Buchan et al., 2014; Bhatia et al., 2021; Delpech et al., 2021; Thiele et al., 2022; White et al., 2025). In contrast, members from the *Fluviicola* genus have been previously associated with increasing temperature and described as late-stage bloom specialists (Delpech et al., 2021), more consistent with predictions that they would be associated with regions with no upwelling. Overall, this emphasizes that microbial community composition patterns are less predictable than those observed for the phytoplankton community, highlighting the need for further work to better resolve the factors governing species-level niche preferences in these complex systems.

An additional influence that glaciers have on the marine microbial community is through the seeding of the marine environment with taxa from supraglacial and/or sub-glacial ecosystems, as well as glacier ice and meltwater rivers has been previously documented (Garcia-Lopez et al., 2019; Thomas et al., 2020). The identification of a *GW2011_GWC1_47_15* (Woesearchaeales) ASV associated at sites without active glacially-induced upwelling may represent an instance of such seeding. This ASV has only been reported in peat bogs or groundwater, both anoxic environments (Groult et al., 2023; Lauzon et al., 2024; Patel et al., 2024). These taxa have the capacity for high molecular weight organic matter degradation (Juottonen et al., 2020), and more broadly, taxa from the Woesearchaeales order have been suggested to form syntrophic relationships with methanogens (Boyd et al., 2010; Liu et al., 2018), a strategy which may allow for their survival in subglacial ecosystems. Another likely example of meltwater-mediated delivery of freshwater taxa to the ocean is the observation of *Tychonema* (Cyanobacteria) in active upwelling regions. *Tychonema* is a mat-forming prokaryote, and been recorded in polar stream and soil communities, as well as alpine lakes and rivers (de los Ríos et al., 2015; Salmaso et al., 2016; Krauze et al., 2021; Pessi et al., 2023; Jablonska et al., 2024). In this study, we observed *Tychonema* exclusively at Sydkap (2022), indicating that its presence reflects a local influence. The Sydkap site in 2022 was characterized high levels of turbidity, likely due to meltwater inputs from the eastern marginal stream and subglacial discharge, making it plausible that *Tychonema* originated from a freshwater environment. Although these freshwater-derived taxa were present during the sampling period, increased sampling resolution throughout the melt season is required to determine whether they persist following water-column mixing.

### Delivery of deepwater taxa to the surface ocean through glacier-induced upwelling

In this study both *SUP05* and *SAR11* exhibited increased relative abundances in deep water, and ASVs belonging to these genera were identified as being associated with regions with active upwelling. Given the occurrence of *SAR11* and *SUP05* in the deeper waters in this study, and previous observations of them throughout the water column, spanning from anoxic to oxic conditions, we expected that these taxa would also occur in surface waters in our study region (Kraemer et al., 2020; Morris & Spietz, 2022; Hays & Fuchsman, 2025). However, their enrichment in deeper waters suggests either fine-scale niche partitioning driven by greater availability of sinking, phytoplankton-derived organic compounds (e.g., osmolytes and methylated amines), or that the lineages detected here represent deep water representatives (Grote et al., 2012; Kraemer et al., 2020; Morris & Spietz, 2022; Hays & Fuchsman, 2025).

The observation of *Cand. Nitrosopumilus* – important marine nitrifiers – in the upper water column in this study is consistent with the vertical advection of deep-water taxa to surface waters along with nutrient-rich ambient waters. In the Arctic, *Cand. Nitrospumilus* is typically observed in surface waters during the winter, where the sun remains below the horizon for months at a time and persistent sea ice coverage limits light penetration, allowing for full water column mixing and nutrient renewal (Christman et al., 2011; Alonso-Sáez et al., 2012; Thiele et al., 2022; Thiele et al., 2023). The presence of *Cand. Nitrosopumilus* in the upper water column presents an intriguing question: are these taxa actively conducting nitrification while at the surface, and if so, how does this influence local nitrogen cycling? Active surface nitrification would imply in situ NO_3_^-^production, potentially resulting in a shift in the surface ocean NO_3_^-^ budget with implications for phytoplankton productivity, and by extension, microbial community structure and ecosystem function. While previous work has identified low potential nitrification rates and limited expression of nitrification genes in Arctic surface waters (Christman et al., 2011; Griffin et al., 2025; Laso-Pérez et al., 2025), we observed that nitrifiers were largely absent in surface waters at sites with no active upwelling. Together, these findings suggest that glacially-induced upwelling exerts a broader influence on marine nitrogen cycling than previously recognized.

Not all instances of deep taxa in the surface waters may be attributed to glacially-induced upwelling. Similar to sites influenced by active glacially-driven upwelling, stations from the Jakeman, Starnes.1, and Starnes.2 sites were characterized by SAR11, with the Jakeman station also exhibiting *SUP05* and *Cand. Nitrosopumilus*, despite being classified as no-upwelling sites. Although bathymetric data are limited, available information (not shown) indicates the presence of a slope on the ocean floor approximately 300 m from the nearest stations sampled along the Jakeman and Starnes transects. We therefore hypothesize that in these instances, the presence of these deep taxa in surface waters reflects delivery via topography-induced upwelling, rather than meltwater-driven upwelling from tidewater glaciers. Additional physical mechanisms, including tidal mixing over sills and wind mixing, could also drive vertical upwelling of deep waters at any of the sites studied. However, physical and geochemical signatures at “Upwelling” classified sites, such as turbidity inputs and isopycnal sloping near the glacier face, indicate that the dominant mode of mixing at the time of sampling is most likely glacially-driven. Further, the disparity in physical and geochemical characteristics between both the glacially-driven and topographically-drive (Jakeman, Starnes.1, Starnes.2) upwelling sites and the remaining non-upwelling sites suggests that the impact of these other mechanisms of physical mixing on the water column were not pronounced.

### Limitations and future directions

It is important to recognize that this study captures only a narrow temporal window. The data collected reflect conditions during a discrete period each year, typically late in the melt season, and cannot resolve finer-scale variability within the season or across years. Logistical barriers are a major cause for this sampling limitation, as access to polar field sites is inconsistent, weather delays are routine, and most research efforts end up biased towards short summer campaigns. Consequently, many key environmental transitions – especially those occurring during winter, spring onset, and fall mixing – remain undersampled. Higher resolution sampling across the melt season, coupled with functional approaches such as metatranscriptomics and metaproteomics, is essential to better understand how ecosystems influenced by glacially-driven upwelling shift over time and respond to change. Realistically, consistent year-round sampling in these regions is only achievable through collaboration with Inuit communities, whose on-the-land expertise and access make this feasible. Collaborative partnerships with Inuit communities, which ensure that research priorities are rooted in community-driven questions, should thus be a key focus of future Arctic research programmes.

## Conclusions

Through a comparative analysis of sites influenced by active tidewater glacier-induced upwelling versus those without upwelling, we identified significant differences in the surface microbial community structure. Upwelling regions were characterized by the presence of deep taxa in surface waters, likely transported upward with nutrient-rich deepwater. Their presence indicates the potential for distinct biogeochemical cycling in these regions, such as altered nitrogen cycling or enhanced carbon cycling, relative to regions lacking upwelling. We also detected taxa associated with upwelling environments that resemble the copiotrophic groups observed during phytoplankton blooms in other polar systems, suggesting that elevated organic matter supply indirectly shapes surface microbial assemblages. At the same time, the presence of taxa from the same genera associated with both upwelling and non-upwelling environments points to fine-scale niche differentiation and species level habitat partitioning that requires further investigation. Together, these results imply that continued glacial retreat, and the resulting shifts in the glacially-driven upwelling regime at sites with tidewater glaciers, may drive a shift towards microbial communities adapted for more oligotrophic conditions, with consequences for nitrogen and carbon cycling. Future efforts should seek to quantify the metabolic activity and productivity of upwelled taxa to clarify their specific biogeochemical roles. More broadly, these findings reinforce the need for higher-resolution temporal sampling to capture the dynamic nature that defines Arctic microbial systems.

## Data Accessibility

Data used in this study is freely and openly available. CTD and biogeochemical data can be found on Canadian Integrated Ocean Observing system at http://doi.org/10.71708/41473632-6b8c-4e8a. DNA sequences were accessioned at NCBI (PRJNA905107). All scripts and associated metadata for analysis can be found on the Bertrand Lab GitHub (https://github.com/bertrand-lab/CAA_16S-analysis).

## Acknowledgements

This work was conducted under licences from the Nunavut Research Institute (2 049 19N-M, 02 030 21R-M). We thank Jimmy Qaapik, the Hamlet of Ausuittuq (Grise Fiord) and the Iviq Hunters and Trappers Association for invaluable consultations, feedback and support. We thank Eric Brossier, France Pinczon du Sel, and Léonie Brossier (*S/Y Vagabond*), Didier Lab members (Laboratoire d’études des littoraux nordiques et arctiques), and the Polar Continental Shelf Project (PCSP) for invaluable assistance in the field. The authors acknowledge financial support from: University of Alberta Northern Research Award, Northern Scientific Training Program, Alberta Graduate Excellence Scholarship funding to JS Spence; Natural Sciences and Engineering Research Council of Canada (NSERC) funding to JS Spence, MP Bhatia, EM Bertrand, D Didier, KO Konhauser, S Waterman; NFRF Explorations Fund grant NFRFE-2018-01,427 to EM Bertrand, S Waterman, MP Bhatia, and J Qaapik; NSERC Shiptime grant to MP Bhatia, S Waterman and EM Bertrand; PCSP grant to MP Bhatia; Canada Research Chairs funding to EM Bertrand and S Waterman; Crown-Indigenous Relations and Northern Affairs Canada funding to D Didier and the Hamlet of Grise Fiord, Campus Alberta Innovation Program Chair funding to MP Bhatia. We thank Martin Kulla for generation of arcGIS figures.

## Supplementary figure and table captions

**Table S1.** Summary of sample metadata and upwelling indicator classifications.

**Figure S1.** Map of stations sampled in 2021 and 2022 spanning A) Harbour Fiord, B) Grise Fiord and in front Grise Fiord community (Ausuittuq), C) Starnes.1 and Starnes.2, D) Jakeman Glacier, E) Sverdrup Glacier, F) Sydkap Glacier and outer South Cape Fiord, G) Belcher Glacier, H) Talbot Inlet, I) Open Jones Sound. Community of Ausuittuq is denoted by the red star. Maps were generated in QGIS as described in Figure 1.

**Figure S2.** Section plots of the upper 75 m of the water column at A) Harbour Fiord, B) Grise Fiord, C) Starnes.1, D) Starnes.2, E) Jakeman Glacier, F) Sverdrup Glacier, G) Sydkap Glacier, H) Belcher Glacier, I) Talbot Inlet. Colour indicates turbidity and density is represented by contour lines. Red triangles indicate stations where bottle samples were collected, and black triangles indicate CTD-only stations. Data is interpolated between stations.

**Figure S3.** As in Figure S2 but for chlorophyll *a* in colour.

**Figure S4.** As in Figure S2 but for dissolved O_2_ in colour.

**Figure S5.** As in Figure S2 but for temperature in colour.

**Figure S6.** As in Figure S2 but for salinity in colour.

**Figure S7.** Principal component analysis (PCA) using a Euclidean distance matrix of physical (strat., turb., temp., diss. O_2_, Chl*a*) and chemical (NO_3_^-^, NO_2_^-^, NH_3_, PO_4_^3-^, SiO_4_ ^4-^) variables of the (A) entire water column (green ellipse = below 50 m, grey ellipse = 50 m and above) and (B) upper 50 m of the water column (blue ellipse = upwelling, upper 50 m within 5 km, purple ellipse = no upwelling, upper 50 m within 5 km). Ellipses represent 95% confidence intervals and are significantly different from one another (PERMANOVA p-value < 0.05). Shapes represent the years sampled, and colours indicate the site sampled. Strat. = stratification; turb. = turbidity; temp. = temperature; diss. O_2_ = dissolved oxygen; Chl*a* = chlorophyll *a*. Percentage values on axis labels represent the total variance in the dataset explained by that principal component.

**Figure S8.** Images taken of Sydkap glacier terminus in South Cape Fiord using Copernicus Sentinel-2A/B data (A) one day before sampling in 2021 (August 13^th^, 2021) (B) sampling date of Sydkap_7_2022 (August 22^nd^, 2022), (C) sampling date of Sydkap_1_2022, and (D) one day before sampling Sydkap_SDB1_2022. Red arrows point to suspected turbidity from river influence.

**Figure S9.** Shannon diversity of upwelling groupings in A) both years, B) 2021, C) 2022.

**Figure S10.** Average relative abundance of microbial taxonomic groups in the upper 50 m of the water column. Genera found at less than 1% average relative abundance were classified as “Other (<1%)”. Samples are grouped according to their Upwelling classification, as indicated by the coloured bar.

**Figure S11.** Microbial community composition utilizing a non-metric multidimensional scaling plot (NMDS) ordination of the (A) entire water column (stress = 0.11; green ellipse = below 50 m, grey ellipse = 50 m and above) and (B) upper 50 m of the water column (stress = 0.18; blue ellipse = upwelling, upper 50 m within 5 km, purple ellipse = no upwelling, upper 50 m within 5 km). (C) Distance-based redundancy analysis (dbRDA) of the microbial community composition in the upper 50 m of the water column with phosphate (PO_4_^3-^), nitrate (NO_3_^-^), ammonia (NH_3_), salinity (Sal.), turbidity (Turb.), temperature (Temp.) and dissolved oxygen (Diss. O_2_ (μM)). Adjusted R^2^ represents the proportion of variation that is explained by the significant environmental predictors (adj. R^2^ = 23%). Percentage value on axis labels represent the proportion of model-explained variance. Vectors indicate significant physical and chemical variables (P < 0.05) as selected by forwards-backwards selection. Shapes represent the years sampled, and colours indicate the site sampled. Ellipses represent 95% confidence intervals and are significantly different from one another (PERMANOVA p-value < 0.05).

**Table S2.** PERMANOVA significance values for all tested combinations of groupings in the upper 50 m and entire water column. * ≤ <0.05; ** ≤ 0.01; *** ≤ 0.001.

